# SpaceSequest: A unified pipeline for spatial transcriptomics data analysis

**DOI:** 10.1101/2025.09.15.676389

**Authors:** Yu H. Sun, Sarbottam Piya, Zhengyu Ouyang, Yirui Chen, Jake Gagnon, Shaolong Cao, Haotian Zhang, Baobao Song, Jing Zhu, Khyati Chandratre, Houlin Yu, Wenxing Hu, Matthew Ryals, Fergal Casey, Dann Huh, Baohong Zhang

**Affiliations:** Research, Biogen Inc., Cambridge, MA, 02142, USA; Data Science, BioInfoRx Inc., Madison, WI, 53719, USA; Biostatistics, Biogen Inc., Cambridge, MA, 02142, USA; PharmaLex, Conshohocken, PA, 19428, USA

**Author notes:** Contribute equally (Yu H. Sun, Sarbottam Piya, Zhengyu Ouyang). Correspondence: Yu H. Sun.

**Keywords:** Spatial transcriptomics, Visium, Visium HD, Xenium, GeoMx, CosMx, single-cell RNA-seq, deconvolution

## Abstract

**Background:** Spatial transcriptomics has emerged as one of the most powerful tools for gaining biological insights, enabling researchers to uncover intricate relationships between gene expression patterns and tissue architecture. Recent advances in the field have resulted in a variety of new platforms, including Visium, Visium HD, and Xenium from 10x Genomics, as well as GeoMx and CosMx from NanoString Technologies, which has now been acquired by the Bruker Corporation. However, the existence of diverse spatial transcriptomics platforms and various data formats poses challenges in standardizing data analysis. Thus, there remains a critical gap in the availability of a comprehensive pipeline capable of conducting end-to-end analysis that is necessary to extract biological insights from multiple spatial transcriptomics platforms.

**Results:** Here, we present SpaceSequest, a tailored pipeline that utilizes cutting-edge computational methodologies to conduct a thorough analysis, enabling the extraction of crucial biological insights from five major spatial transcriptomics technologies. SpaceSequest performs (1) standardized quality control and general data processing, (2) key analyses customized for each spatial platform, (3) automated cell type annotation and deconvolution, and (4) high-quality figure and analysis result generation. In addition, SpaceSequest allows for smooth integration with cellxgene VIP and Quickomics for user-friendly data access and interactive visualization.

**Conclusions:** SpaceSequest is a unified and comprehensive pipeline designed for the analysis, visualization, and publication of spatial transcriptomics data from various platforms. The source code is available at https://github.com/interactivereport/SpaceSequest. To facilitate seamless installation and usage, we have also created a detailed Bookdown tutorial that can be accessed through https://interactivereport.github.io/SpaceSequest/tutorial/docs/index.html.

## BACKGROUND

The advent of single-cell RNA sequencing (scRNA-seq) has revolutionized our understanding of cellular heterogeneity and gene expression dynamics. It represents a significant advancement from bulk RNA-seq, which only provides average expression profiles from mixed cell populations, thereby obscuring the underlying cellular diversity [1–3]. Although scRNA-seq generates invaluable insights into cellular heterogeneity, it lacks spatial information, as it dissociates cells from their original tissue context [4, 5]. Understanding the intricate spatial organization of gene expression within complex tissues and organs is crucial for unraveling the mysteries of biology [6–8]. Spatial transcriptomics (ST) has emerged as a transformative technology that quantifies RNA within intact tissue sections, enabling researchers to explore gene expression patterns within their native spatial context [6, 9–12].

ST can provide precise insights into disease progression by discerning heterogeneity within diseased tissue, which is powerful for studying complex diseases such as tumor and neurological disorders[13–17]. For example, using ST data, Berglund et al. not only distinguished healthy and diseased areas but were able to identify re-stratified tumor heterogeneity by analyzing gene expression gradients within the tumor microenvironment, specifically in the stroma adjacent to the tumor area [18]. These advancements have unlocked new avenues for addressing critical biological inquiries. First, ST plays a pivotal role in characterizing the cell-type composition of tissues. While definitions are often derived from large-scale single-cell RNA-seq or epigenetics datasets and projected onto ST data, the technology also empowers the direct generation of cell-type definitions from the spatial transcriptomics data itself[19, 20]. This application has been widely utilized, resulting in the creation of compositional atlases for a diverse range of tissues, including the nervous system [21–24], human kidney [25–27], heart [28, 29], testes [30], and lung [31, 32]. Second, ST facilitates the exploration of cellular interactions, revealing the rules and patterns governing spatial covariation among distinct cell types. For instance, studies examining the mouse visual cortex identified a preferential spatial proximity of inhibitory neuron subtypes, shedding light on their organizational principles[33]. Similar proximity analyses have been instrumental in elucidating gene expression patterns linked to Alzheimer’s disease and in characterizing histopathological responses to traumatic brain injury[34–37]. Finally, ST’s transcriptome-wide data on in situ gene expression empowers the elucidation of molecular interactions between tissue components[7]. Through the identification of ligand-receptor pairs among cell types at varying spatial proximities, we can discern the intricate communication networks that underlie phenomena such as tumor interactions with the microenvironment, immune infiltrates in tissues, and the establishment of developmental gradients[38, 39].

There are two distinct ST methods: (i) sequencing-based ST, and (ii) imaging-based ST. In the sequencing-based ST method, a DNA-barcoded surface is used to capture RNA molecules from the intact tissue section. These captured RNA molecules are then subjected to next-generation sequencing to determine their respective gene identities[6]. By contrast, in imaging-based methods, multiple mRNA transcripts are simultaneously detected within intact tissue using fluorescence in situ hybridization (FISH) or direct in-situ sequencing[6, 12]. The imaging-based methods, such as 10x Genomics Xenium and NanoString CosMx, are suitable for hypothesis testing as these methods allow transcriptomic profiling at single cell resolution albeit for a limited number of target genes. Conversely, sequencing-based methods which produce an unbiased, whole-transcriptome profile, are suitable for hypothesis generation. Typical methods include 10x Genomics Visium, Visium HD, and NanoString GeoMx with sequencing readout. However, transcriptomics profiles obtained from these methods are not directly resolved at single cell resolution.

One critical drawback of current grid-based ST profiling methods (Visium, Slide-seq [38, 40], and HDST [41] is their inability to achieve single-cell resolution. These techniques either combine multiple cells or fractions of multiple cells at each tissue location, limiting the ability to discern individual cell contributions. Nevertheless, this limitation can be addressed through computational integration, leveraging the reference transcriptome signatures of cell types obtained from coupled single-cell RNA sequencing (scRNA-seq) profiles. By integrating ST data with scRNA-seq data, researchers can deconvolute cell type composition and retrieve further information from the spatially resolved gene expression patterns[42, 43].

The landscape of spatial transcriptomics data analysis has seen the development of several data analysis pipelines, including ST Pipeline [44] and Spacemake [45]. ST Pipeline primarily focuses on the essential tools and scripts required to process raw files derived from Visium data in FASTQ format, providing datasets primed for downstream analysis [44]. In contrast, Spacemake represents a more comprehensive solution, offering a wider range of functionalities compared to ST Pipeline. This tool is adept at handling spatial transcriptomic data from diverse platforms, such as Visium, Slide-seq, Seq-scope, etc [45]. Another distinctive feature of Spacemake is its utilization of NovoSpaRc [46] that allows mapping snRNASeq data to spatial coordinates [45]. Recently developed pipelines are focused on specific tasks, for example, PIPEFISH[47] was designed for FISH spatial transcriptomics, ENACT was specifically designed for Visium HD data analysis[48], while Panpipes[49] was created for spatial multi-omics analysis. Although these existing pipelines are useful, their capabilities fall short in extracting comprehensive and meaningful biological insights from diverse spatial transcriptomics datasets.

Here we introduce SpaceSequest, our comprehensive spatial transcriptomics data analysis pipeline tailored for the analysis of several major platforms, spanning 10X Genomics Visium, Visium HD, Xenium, as well as NanoString GeoMx and CosMx platforms. By utilizing cutting-edge computational methodologies, SpaceSequest streamlines a series of essential analyses to extract meaningful biological insights. Compatible with diverse input formats, the pipeline seamlessly navigates through critical steps including clustering, deconvolution, cell type annotation, and differential expression analysis. This integrated approach equips researchers with a powerful toolkit to dissect the spatially defined molecular landscape, facilitating a deeper understanding of complex biological systems and their microenvironmental interactions.

### IMPLEMENTATION

#### General data processing

SpaceSequest supports various input formats and processes the data through a series of automated steps, leveraged by multiple well-designed packages including Seurat 5 [50] and anndata[51]. For sequencing-based strategies (10x Genomics Visium, Visium HD, and NanoString GeoMx with sequencing readouts), FASTQ processing is required before running

SpaceSequest. The input files for running Visium and Visium HD workflows, including fundamental quality metrics and the unique molecular identifiers (UMI) count matrix, can be generated through 10x Genomics Space Ranger software. Since our pipeline does not rely on gene annotation or species-specific details, users are required to accurately select the appropriate species during Space Ranger execution. For GeoMx, the GeoMx NGS Pipeline developed by NanoString is required to convert FASTQ to digital count conversion (DCC) files, which can be further processed by SpaceSequest. In the case of imaging-based technologies such as Xenium and CosMx, files generated by the platform can be readily used for SpaceSequest data processing.

#### Clustering

Clustering is a powerful tool for understanding the structure of the data, as well as the spatial tissue domains. For general clustering of single cell expression in CosMx and Xenium, or bins in Visium HD, we utilized the UMAP dimension reduction followed by the FindNeighbors and FindClusters functions. For Visium technology, we have incorporated two more clustering tools, SpaGCN [52] and BayesSpace [53]. SpaGCN leverages graph convolutional networks to integrate gene expression, spatial location, and histology data, allowing for the identification of spatial domains with distinct expression patterns [52]. In contrast, BayesSpace is a Bayesian statistical tool that leverages prior spatial knowledge to cluster transcriptomically similar spots in proximity [53]. Additionally, BayesSpace offers the capability to enhance resolution by reallocating gene counts from larger spots in array-based data, such as Visium, to more refined sub-spots based on spatial information from nearby spots [53].

#### Cell type annotation

Automated cell type annotation can be achieved through transferring labels from a reference single-cell RNA-seq dataset. For single-cell-level assays, including CosMx and Xenium, we incorporated Azimuth to predict the most likely cell type for each cell[54]. A UMAP plot will be generated after running the pipeline for users to examine the cell type annotation results.

#### Deconvolution

For deconvolution purposes, we have incorporated two state-of-the-art deconvolution tools, Tangram [55] and Cell2location [42]. We selected these two packages for their outstanding performance in a recent benchmarking study[43]. Tangram, a deep-learning framework, excelled in creating comprehensive single-cell-resolution spatial gene-expression maps and integrating histological and anatomical data from the same specimens. It adeptly handled low-quality genes in high-resolution methods, offered single-cell resolution for lower-resolution techniques, and ensured broad genome-wide coverage for targeted approaches[55]. Meanwhile, Cell2location, a Bayesian model, showcased its ability to discern fine-grained cell types within spatial transcriptomic data and constructs detailed cellular maps of diverse tissues. In the first step, Cell2location estimated reference cell type signatures from single-cell and single-nucleus RNA-seq profiles, aligned with gene expression profiles for user-specified cell types and subpopulations. In the subsequent stage, Cell2location utilized these reference signatures in conjunction with one or more spatial transcriptomic datasets to deconstruct mRNA counts at individual spatial locations into their respective reference cell types[42].

#### Ligand receptor interactions

Here, we employ SpaTalk to investigate spatially resolved cell-cell interactions that will enable us to get insights into tissue homeostasis, development, and disease[56]. This tool leverages a graph network along with a knowledge graph to model and evaluate the ligand-receptor-target signaling network operating between neighboring cells[56]. SpaTalk serves as an optional step in the Visium data processing workflow, providing researchers with a powerful means to unravel the complex dynamics of inter-cellular interactions within tissues[56].

#### Standardized output

SpaceSequest generates a series of output files that are ready for result query and further analysis. The R objects are saved as RData files, and the python anndata objects are saved as h5ad files. These files contain expression matrices, dimension reduction results, and intermediate data generated during pipeline processing.

## RESULTS

### Highlights of SpaceSequest

The pipeline incorporates several key workflows to perform various analyses corresponding to each platform. Key strengths of this pipeline include:

- 1) Self-explainable configuration files that ensure easy set up.
- 2) Standardized and reproducible data processing powered by R and Python.
- 3) Customized tools are integrated in the pipeline for each spatial platform.
- 4) Comprehensive result generation, including R/Python objects and key figures.
- 5) Streamlined data visualization enabled by cellxgene VIP and Quickomics.

### Overall pipeline design

SpaceSequest is a unified toolkit which contains a set of workflows. Each workflow can be activated by calling the workflow script, and necessary configuration files will be created accordingly. The main configuration files are in YAML format, ensuring standardized formatting and flexible modification. To execute the pipeline, in most of the times, users are typically required to fill in a sample metadata file, which documents the details of each data, such as sample name, file path, and associated sample-level information. The overall pipeline components are illustrated in Figure 1.

**Figure 1.**
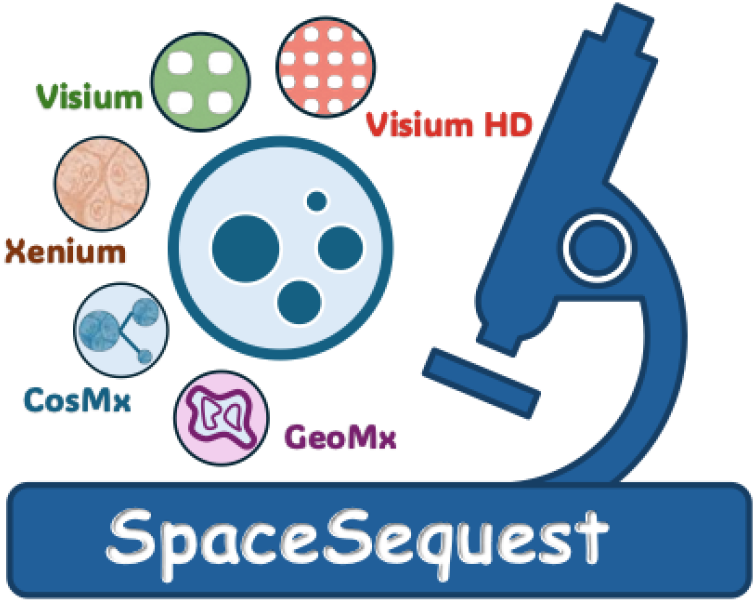
Components of SpaceSequest. The pipeline integrates five major workflows tailored to perform a series of analysis for each platform, including Visium, Visium HD, Xenium, CosMx, and GeoMx.

### Visium data analysis incorporates multiple tools

The 10x Genomics Visium platform was released in 2019 and continues to serve as a powerful commercial spatial transcriptomics technology since then. The Visium slides contain an array of barcoded 55 µm spots, with two major tissue capture sizes: 6.5×6.5 mm or 11×11 mm. Sequencing data in FASTQ format are processed through Space Ranger, and SpaceSequest continues the analysis by utilizing the binned output files. Firstly, SpaceSequest performs standard quality control on the data by removing spots with low gene detection rates, low UMI counts, etc. The filtered data are then integrated into a new h5ad file, which is piped to downstream tools for further analysis.

We tested our pipeline on a public dataset generated from the brain tissue of an Alzheimer’s disease mouse model, which has mutations in APP and PSEN1 genes[57]. We also downloaded the Azimuth mouse motor cortex single-nucleus RNA-seq data as a reference for cell type annotation[58]. The data analysis is achieved by calling the ***visium*** command in SpaceSequest, which reads all input data and performs a series of quality control steps (Figure 2A). The cleaned data are then integrated for downstream processing, which includes spatial clustering and cell type deconvolution results. To leverage the power of parallel computing, SpaceSequest is able to submit sbatch jobs to execute the following four tools simultaneously: BayesSpace[53], Cell2location[42], SpaGCN[52], and Tangram[55]. The jobs will be monitored until results have been generated and saved in separate directories. To provide a few examples, BayesSpace clustering step will cluster the spots across samples and identify comparable spatial clusters (Figure 2B). While Cell2location results can be visualized by cell abundance heatmaps overlayed with the tissue image (Figure 2C). Additionally, the pipeline utilizes SpaTalk for cell-cell communication analysis, which requires Cell2location output so it is only executed once Cell2location results are available (Figure S1)[56]. Overall, our Visium data analysis workflow ensures automated data processing and analysis with a streamlined, one-stop-shop design (Figure 2A).

**Figure 2.**
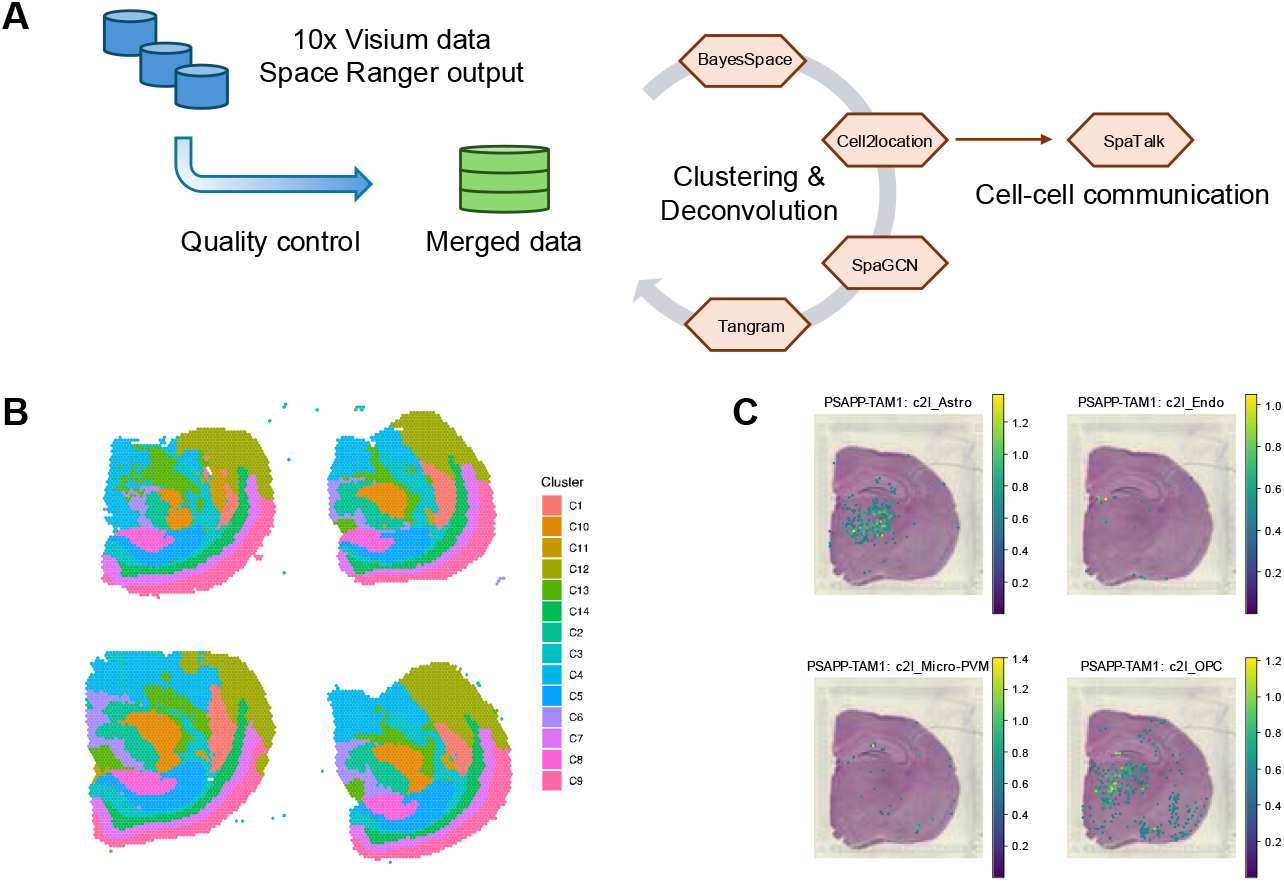
10x Visium data analysis. (A) Workflow for Visium data processing. (B) Spatial clustering results generated by BayesSpace. This run contains four samples, and BayesSpace identified shared clusters across them. (C) Predicted cell type abundance by Cell2location in one sample (PSAPP-TAM1), highlighting astrocytes, OPCs, microglia, and endothelial cells.

### Streamlined Xenium and CosMx data analysis

The 10x Genomics Xenium and NanoString (now acquired by the Bruker Corporation) CosMx platforms are both imaging-based, high-plex in-situ technologies capable of capturing up to thousands of RNAs simultaneously at single-cell resolution. They require a set of probes, usually from pre-designed panels, with additional space for customization as well. Xenium data consists of a set of well-defined files in various formats, such as the cell_feature_matrix files (a folder with three associated files, or a single h5 file), transcript files (transcripts.csv.gz, transcripts.parquet, transcripts.zarr.zip), and cell summary files (cells.csv.gz, cells.parquet), etc. The LoadXenium function from the Seurat package reads the necessary files from the directory and converts them into a Seurat object for downstream analysis.

CosMx data are stored as a collection of plain text files in the comma-separated values (csv) format. These files include a metadata file (metadata_file), a file storing field of view (fov) positions (fov_positions_file), an expression matrix file (exprMat_file), a polygon file (polygons), and a transcript data file (tx_file). Data loading is achieved by the LoadNanostring function, which converts these csv files to a Seurat object.

SpaceSequest contains two workflows—***xenium*** and ***cosmx*** —to streamline the process from data loading to downstream analysis. Once Xenium or CosMx data are successfully loaded, these workflows incorporate standard procedures that perform quality control, clustering, cell type annotation, and differential expression (DE) analysis, etc (Figure 3A). Moreover, the workflows generate UMAP and tissue visualizations, enabling users to examine the dimension reduction and cell type annotation results. Example plots generated using public datasets are shown in Figure 3B-D. The Xenium data were generated from a healthy human brain FFPE sample profiled with the Xenium Human Brain Gene Expression Panel containing 266 targets. On the other hand, the CosMx data were derived from human frontal cortex FFPE tissue using >6,000 target genes. After running the workflow, various analysis results and the processed Seurat object will be saved for future queries and additional analyses, including further exploration by Cellxgene VIP[59].

**Figure 3.**
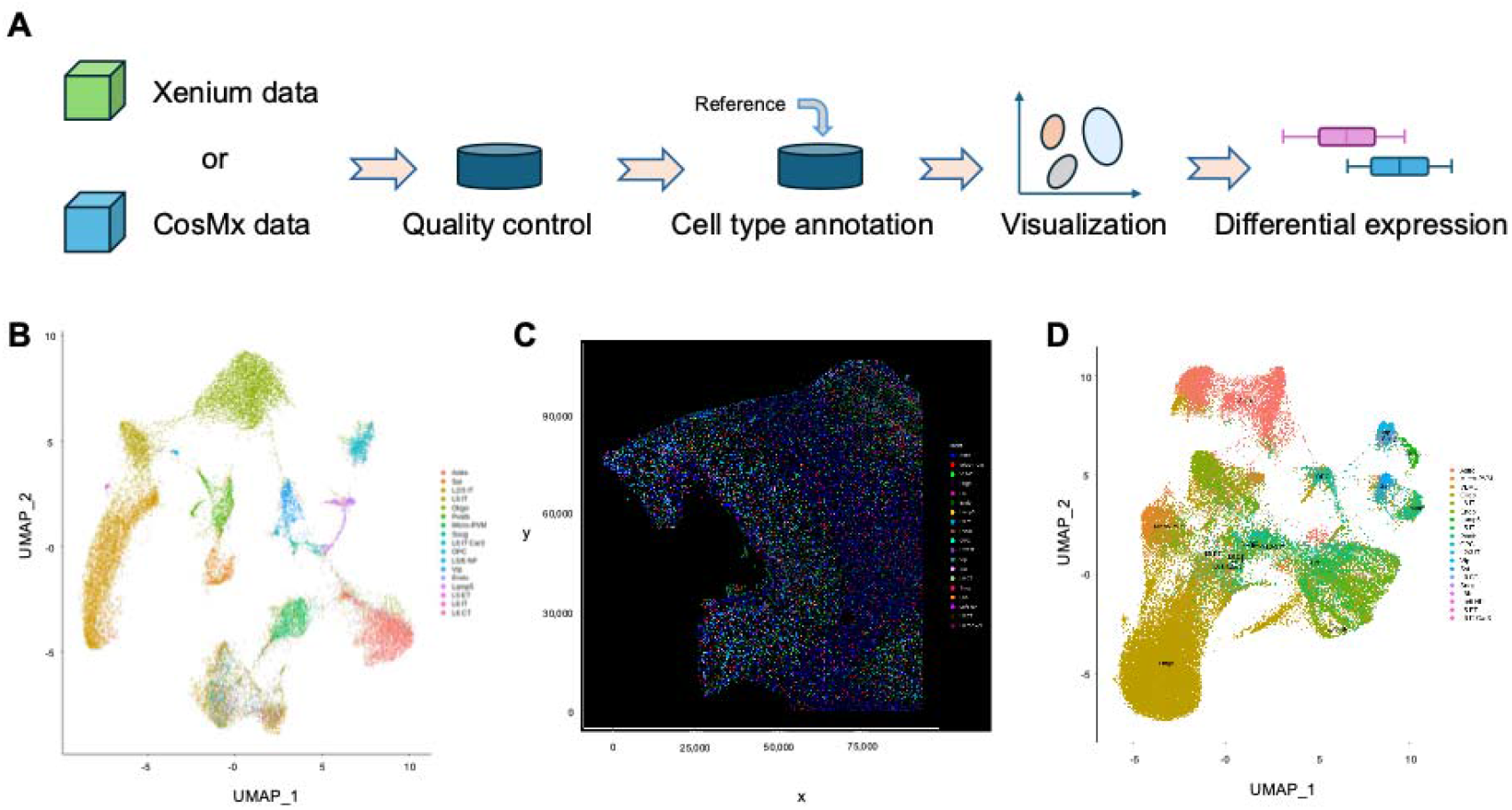
Xenium and CosMx data analysis. (A) Streamlined workflows for analyzing Xenium and CosMx data. (B) UMAP visualization of Xenium data, labelled by predicted cell types. (C) Tissue and cell visualization of CosMx data. (D) UMAP visualization of CosMx data with predict cell types labelled.

### Enhanced GeoMx data processing and visualization

The NanoString GeoMx platform is a useful technique to compare gene expression differences in defined regions on the tissue slides. It uses probes to detect the whole transcriptome in the selected region of interests (ROIs). However, GeoMx is not a single-cell resolution technique; instead, it requires a sufficient number of cells within the region to generate robust signals. Thus, GeoMx is considered as a mini-bulk RNA detection platform, capable of profiling >18,000 major genes for human and mouse samples. In our pipeline, we developed ***geomx***, a workflow that takes DCC files, sample metadata, and a set of comparison groups as input. It performs a series of analysis, and outputs differential expression analysis results (Figure 4A). This workflow generates four csv files that are fully compatible with other tools including some Shiny applications[60, 61], for example, Quickomics, an open-source R Shiny application for flexible data exploration and visualization[60]. After uploading the data, users can examine PCA plots (Figure 4B), volcano plots (Figure 4C), heatmaps (Figure 4D), and a number of on-the-fly visualizations through Quickomics. This strategy not only automates GeoMx data analysis, especially for studies with many samples, but bridges the gap between raw data and insight generation through Quickomics, which creates a link to facilitate data storage and future analysis.

**Figure 4.**
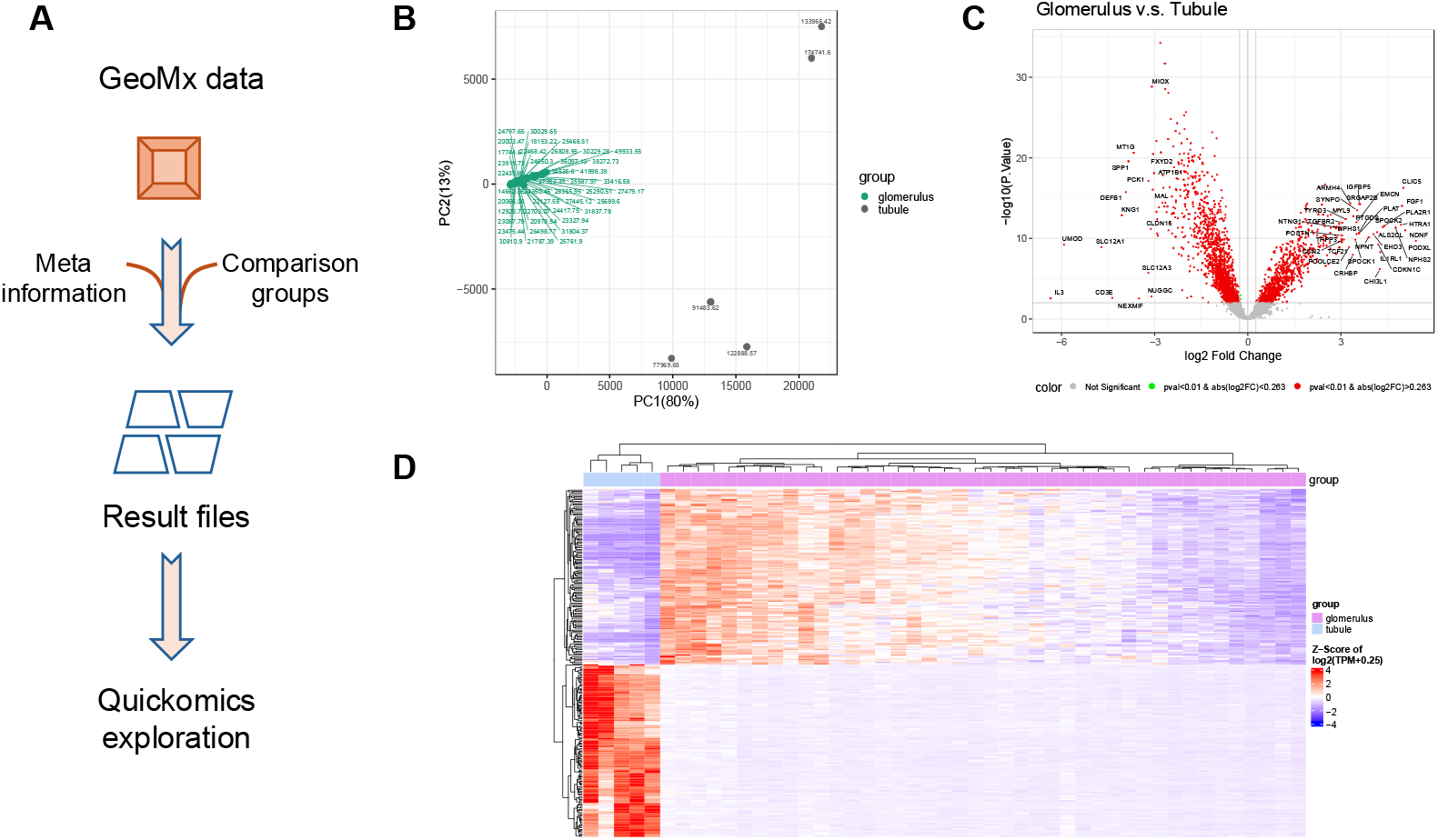
GeoMx data analysis. (A) Overview of GeoMx data analysis workflow. (B) PCA plot showing samples from two kidney regions, glomerulus and tubule. (C) Volcano plot highlighting differentially expressed genes between glomerulus and tubule. (D) Heatmap displaying the top differentially expressed genes between glomerulus and tubule, with samples and genes clustered by dendrograms.

### Standardized data analysis of the cutting-edge Visium HD platform

In 2024, 10x Genomics released a new platform called Visium HD, featuring high-resolution, gap-free detection of tissue RNAs. This platform is now compatible with FFPE, fixed frozen, and fresh frozen samples. To keep pace with the rapid evolution of spatial transcriptomics technologies, we incorporated ***visiumhd*** into SpaceSequest for Visium HD data processing. The workflow takes Space Ranger outputs, performs bin-level quality control, dimension reduction, and cell type annotation when an Azimuth reference dataset is available (Figure 5A). By executing the pipeline, users receive plots to examine the gene counts (Figure 5B) and cell type annotation results (Figure 5C). Furthermore, we developed a new function, getVisiumHDspotCol, to extract the mean RGB intensities when a paired high-resolution immunofluorescence image is available (Figure 5D). This function facilitates accurate bin selection using nuclei staining, cell type marker staining, or other biomarker staining, for example, pathological protein markers.

**Figure 5.**
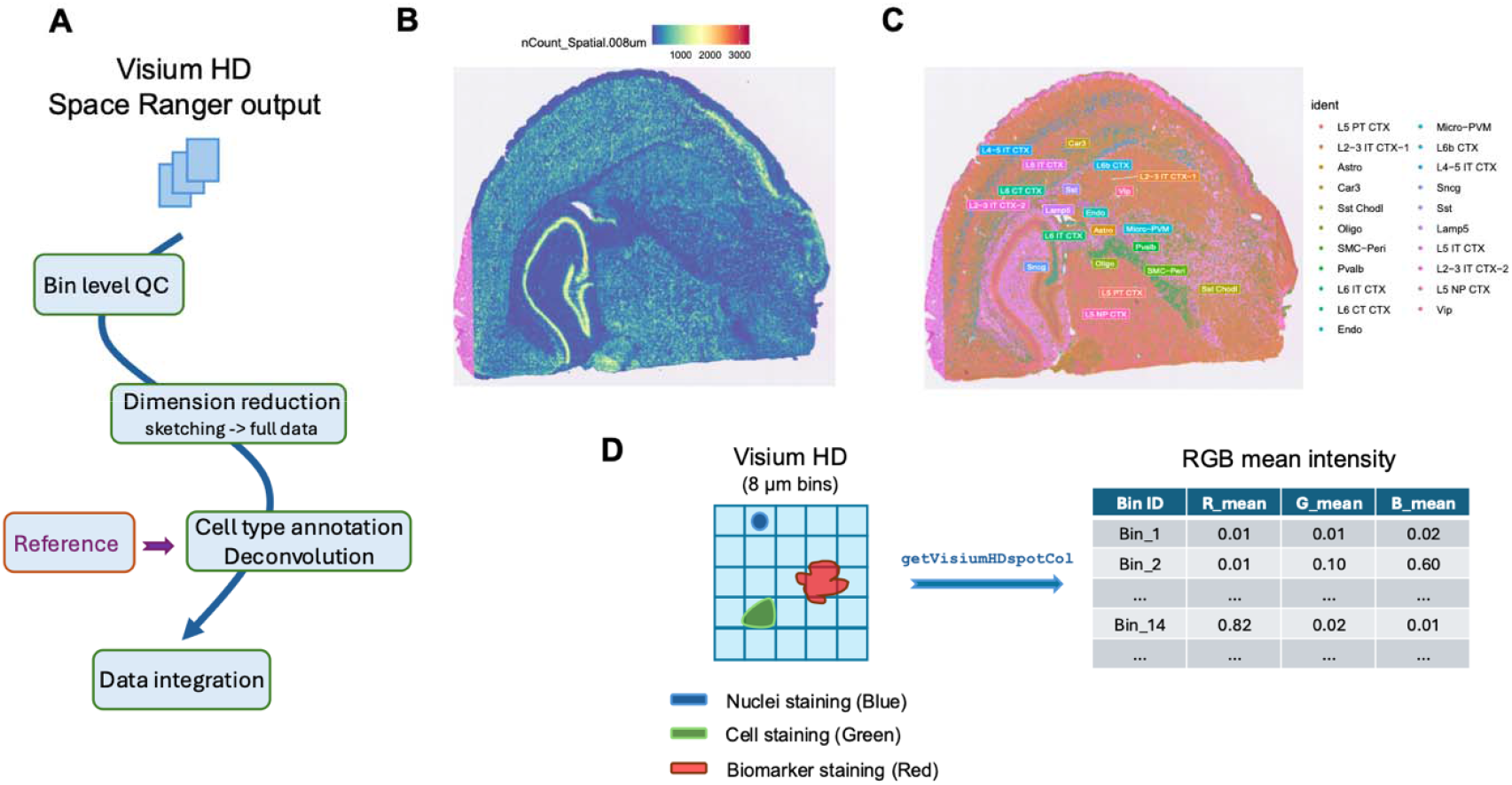
Visium HD data analysis. (A) Overview of the general processing workflow for Visium HD data. (B) Heatmap showing the number of UMI counts (nCount) detected across the tissue section at 8 μm bin resolution. (C) Visualization of annotated cell types on the tissue section. (D) Illustration of a new function that calculates the average RGB intensities across bins.

## DISCUSSION

In this study, we developed SpaceSequest, a unified pipeline for spatial transcriptomics data processing. Currently, it supports five major platforms and incorporates customized software and packages to maximize the information that can be extracted from the data. To make the pipeline accessible, we provided a detailed tutorial aimed at lowering the barrier for researchers to apply this SpaceSequest to their own studies. Guided by the FAIR principles (Findable, Accessible, Interoperable, and Reusable), SpaceSequest enhances data access, interpretation, and facilitates data sharing by generating standardized results. In summary, SpaceSequest is a user-friendly, end-to-end solution for spatial transcriptomics data analysis and interpretation.

## Abbreviations

scRNA-seq: Single-cell RNA-seq
NGS: Next-generation sequencing
csv: comma-separated values
fov: field of view
ST: spatial transcriptomics
UMI: unique molecular identifiers
FISH: fluorescence in situ hybridization
DE: Differential expression
DCC: digital count conversion
PCA: Principal component analysis
OPC: Oligodendrocyte progenitor cells
UMAP: Uniform Manifold Approximation and Projection
Cellxgene VIP: Cellxgene visualization in plug-in
LRI: ligand-receptor interaction

## Ethics approval and consent to participate

Not applicable.

## Consent for publication

Not applicable.

## Author Contributions Statement

Y.H.S., S.P., and Z.O. developed the pipeline and wrote the manuscript, with the help from Y.C., J.G., S.C., H.Z., B.S., J.Z., K.C., H.Y., W.H., M.R., F.C., D.H., and B.Z. All authors reviewed the manuscript.

## Acknowledgement

We acknowledge the scientists at Biogen for their valuable feedback on the pipeline.

## Availability and Requirements

No new data were generated from this study. Several public datasets were used to demonstrate SpaceSequest. The 10x Visium data were from a public study using mouse brain tissues with NCBI accession number GSE203424 (https://www.ncbi.nlm.nih.gov/geo/query/acc.cgi?acc=GSE203424). Public data were downloaded from 10x Genomics dataset webpage (https://www.10xgenomics.com/datasets) to test Xenium (https://www.10xgenomics.com/datasets/xenium-human-brain-preview-data-1-standard) and Visium HD (FFPE: https://www.10xgenomics.com/datasets/visium-hd-cytassist-gene-expression-libraries-of-mouse-brain-he, Fixed frozen: https://www.10xgenomics.com/datasets/visium-hd-cytassist-gene-expression-mouse-brain-fixed-frozen) workflows, respectively. Moreover, we used a public human kidney GeoMx dataset (http://nanostring-public-share.s3-website-us-west-2.amazonaws.com/GeoScriptHub/Kidney_Dataset_for_GeomxTools.zip) and a human brain CosMx dataset (https://nanostring.com/products/cosmx-spatial-molecular-imager/ffpe-dataset/human-frontal-cortex-ffpe-dataset/) from NanoString websites to test GeoMx and CosMx pipelines respectively. More details related to data downloading and processing can be found at the tutorial (https://interactivereport.github.io/SpaceSequest/tutorial/docs/index.html) associated with this pipeline.

Project name: SpaceSequest

Project home page: https://github.com/interactivereport/spacesequest

Operating system(s): Linux, Mac OS

Programming language: Bash (Unix Shell)

Other requirements: Conda

License: MIT license

Any restrictions to use by non-academics: No

## Supplementary figure legends

Figure S1: Examples of SpaTalk results. (A) Highlighted pairs between astrocytes (Astro) and GABAergic neurons. (B) Significantly enriched ligand-receptor interaction (LRI) pairs between astrocytes and GABAergic neurons. (C) Highlighted pairs between microglia and macrophage cells (Micro_PVM) and Glutamatergic neurons. (D) Significantly enriched ligand-receptor interaction (LRI) pairs between microglia/macrophages and Glutamatergic neurons.

