## Supplementary figures and images for "SpaceSequest: A unified pipeline for spatial transcriptomics data analysis"

### Supplementary figure

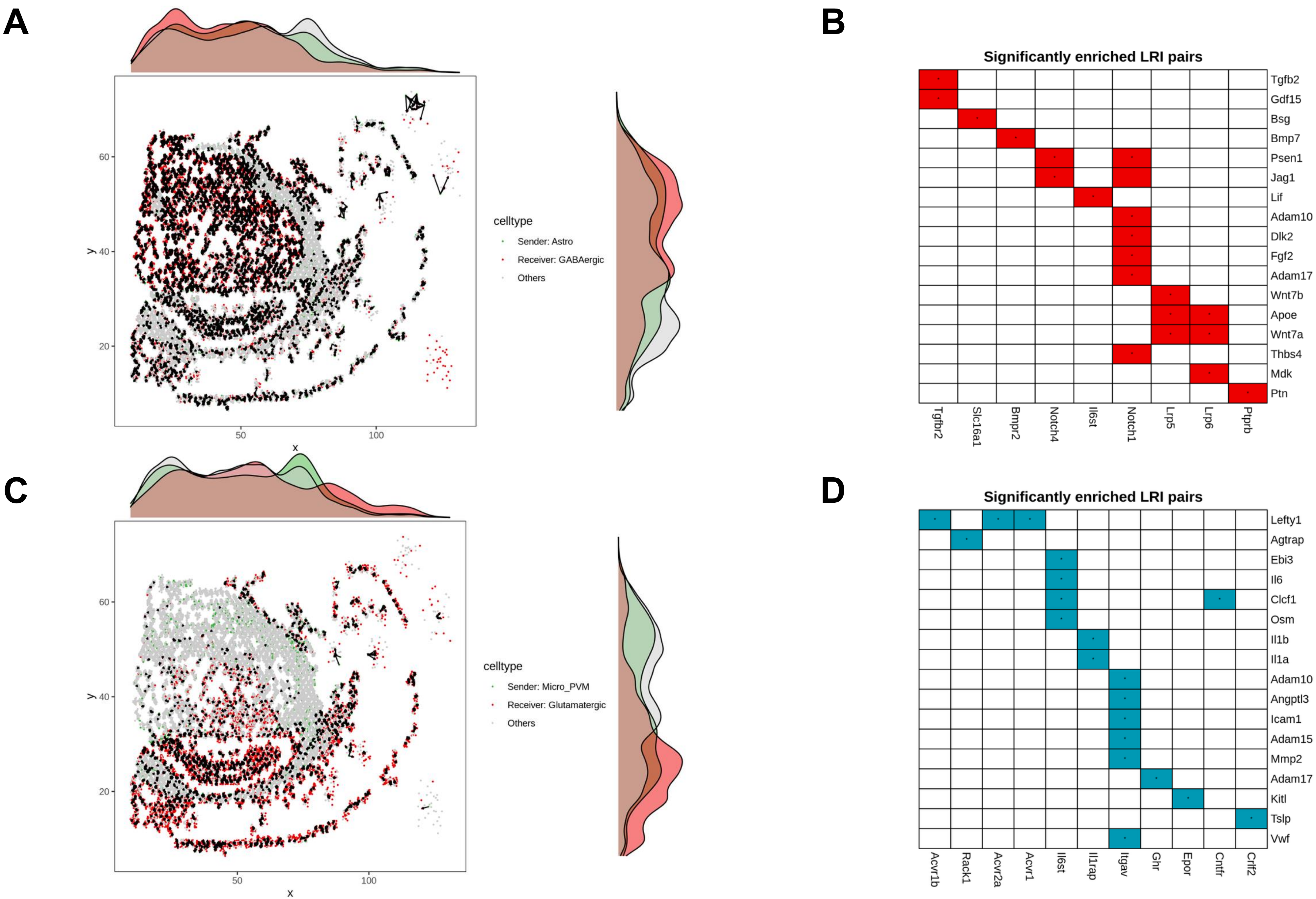
